# Establishing *de novo* microbial biosynthesis of anisole

**DOI:** 10.64898/2026.11.02.763652

**Authors:** Lyon Bruinsma, Diego Rosas Gomez, Costanza La Rocca, Jose Eduardo Escrig Molina, Vitor A. P. Martins dos Santos, Christos Batianis

**Affiliations:** Bioprocess Engineering, Wageningen University & Research, Wageningen, 6708 WE, The Netherlands; LifeGlimmer GmbH, Berlin, 12163, Germany

**Keywords:** anisole, *Pseudomonas putida*, *O*-methyltransferase, growth-coupled selection, directed evolution, *de novo* biosynthesis, two-phase extraction, shikimate pathway

## Abstract

Anisole is a valuable aromatic ether and promising green solvent whose industrial manufacture relies exclusively on petroleum-derived phenol and hazardous alkali-mediated methylation. Here, we establish the first *de novo* microbial biosynthesis of anisole from glucose in *Pseudomonas putida* KT2440 by integrating retrosynthetic pathway design with growth-coupled directed evolution. Using a selection platform in an engineered double auxotroph, in which growth depends on S-adenosylmethionine-dependent methylation, we evolved a promiscuous plant chavicol O-methyltransferase to yield phenol O-methyltransferase 1 (POMT1; bearing a key D264N substitution), which improved anisole production by 2.3-fold over the wild-type enzyme. To overcome product volatility, we integrated an *in situ* two-phase extraction system using dodecane that achieved near-quantitative product recovery. Then, we compared two phenol routes from the shikimate pathway, selected the 4-hydroxybenzoate route, and raised precursor supply by deleting competing reactions and introducing feedback-resistant and reduced-flux variants in two shikimate pathway enzymes. The final strain produced up to 0.57 mM anisole (62 mg/L) from glucose.

## INTRODUCTION

Anisole (methoxybenzene) is a structurally simple, yet highly valuable aromatic ether widely used in the cosmetics, pharmaceutical, agrochemical industries and food industry, where it is used as a flavoring agent, due to its characteristic sweet, phenolic and anise-like aroma.^1^ Although it occurs naturally in trace quantities in certain plant essential oils,^2^ direct extraction from biomass is economically unviable,^3^ and industrial supply is therefore met exclusively through petroleum-based chemical synthesis, principally the liquid-phase methylation of phenol with hazardous alkali reagents.^1^ Gas-phase catalytic methylation routes exist as alternatives, but they require high operating temperatures, suffer from catalyst deactivation and generate significant impurities.^4^ Additionally, every existing route still depends on fossil-derived phenol, which keeps anisole production disconnected from renewable feedstocks, contrary to the aims of green chemistry.^5,6^

Microbial cell factories offer a compelling alternative for the sustainable synthesis of industrial chemicals, combining high selectivity with mild, aqueous operating conditions.^7,8^ Hosts such as *Pseudomonas putida*,^9^ *Escherichia coli*^10^ and *Saccharomyces cerevisiae*^11^ have already been engineered to produce a broad array of non-native compounds, so the binding constraint is no longer the host but the pathway. However, many industrially relevant molecules, including anisole, have no described biosynthetic route in a microbial host. Retrosynthetic algorithms address this limitation by mining enzymatic reaction rules to predict new pathways, often by exploiting the latent promiscuity of existing enzymes to build “underground” metabolic routes to new-to-nature compounds.^12,13^ However, the non-native activities predicted *in silico* typically show suboptimal kinetics, and improving them by laboratory evolution has been constrained by the need for a high-throughput assay, since microplate-based screening of large mutant libraries is labor-intensive. A selection that links cellular growth to the non-native activity would remove this constraint, because improved variants could be enriched by growth instead of being identified one by one. Here, we report the complete *de novo* biosynthesis of anisole from glucose in an engineered *P. putida* KT2440. First, we used retrobiosynthetic selection to identify a promiscuous plant *O*-methyltransferase with activity on phenol and, due to anisole’s volatility, implemented an *in situ* biphasic extraction platform with a dodecane overlay to retain the product during cultivation. Then, we built a growth-coupled selection system to improve this enzyme and isolated a variant that enhanced anisole production by 2.3-fold relative to the wild-type enzyme. Finally, we engineered upstream precursor supply by comparing two decarboxylation routes to phenol and increasing carbon flux through engineering the shikimate and pentose phosphate pathways, reaching a *de novo* anisole titer of 0.57 mM (62 mg/L).

## RESULTS AND DISCUSSION

### Retrosynthetic Identification of a Phenol *O*-Methyltransferase

The biological synthesis of anisole requires the *O*-methylation of phenol, a reaction documented to be catalyzed by phenol *O*-methyltransferase (EC 2.1.1.25). Although this enzymatic activity has been identified in mammals, no coding sequence is currently available to enable heterologous expression in microbial hosts.^14^ We therefore used Selenzyme, an enzyme selection tool for pathway design, to search for promiscuous methyltransferases whose substrate specificities are structurally analogous to phenol (Table S4).^15,16^

The search prioritized CVOMT1 (Q93WU3), a chavicol *O*-methyltransferase from *Ocimum basilicum* (EC 2.1.1.146) that natively transfers a methyl group from S-adenosyl-L-methionine (SAM) to chavicol to produce estragole.^17^ An earlier *in vitro* study had already shown that CVOMT1 also accepts phenol, with modest activity (10.9% of its activity on chavicol), yet anisole formation had not been reported.^17^ Because CVOMT1 and phenol *O*-methyltransferase share the same reaction chemistry and cofactor, we reasoned that anisole should be the product of this promiscuous activity.

### Host Tolerance and In Situ Two-Phase Extraction for Anisole Retention

Before testing CVOMT1 in *P. putida* KT2440, we first evaluated host tolerance to the substrate phenol, as aromatic compounds, particularly phenolic intermediates, frequently exhibit cellular toxicity toward bacterial hosts.^18^ Growth of *P. putida* in minimal medium with glucose across 0 to 5 mM phenol showed that specific growth rates remained unaffected up to 2 mM (0.78 ± 0.02 h ¹) and decreased moderately to 0.58 ± 0.02 h ¹ at 5 mM (Figure S1A), consistent with the membrane-disruptive effects of phenolic hydroxyl groups.^19–22^

Next, we evaluated product toxicity across the same concentration range, and no growth inhibition was observed at any tested anisole concentration (Figure S1B). However, anisole is highly volatile, with 89% lost by evaporation from a 2 mM cell-free aqueous solution within 16 h (Figure 1A). Consequently, this rapid loss prevents definitive conclusions regarding cellular tolerance, as the effective exposure concentration was likely substantially lower than nominal levels. Furthermore, volatilization would lead to product loss in standard cultivation setups, highlighting the necessity of an *in situ* two-phase extraction strategy.

**Figure 1.**
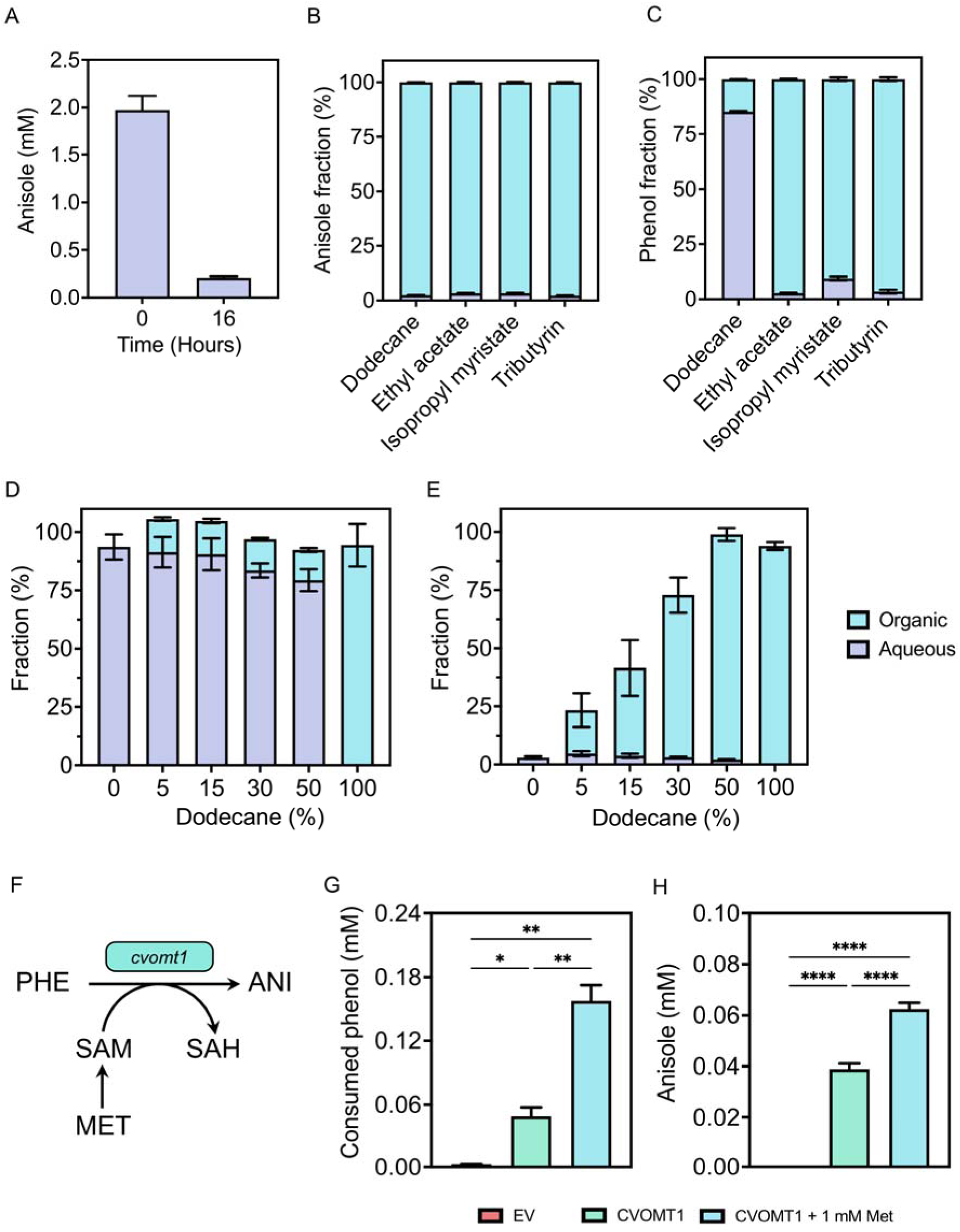
*In situ* product capture and *in vivo* demonstration of anisole production by CVOMT1. (A) Abiotic evaporation of anisole over time. (B) Anisole partition in selected organic solvents. (C) Phenol partition in selected organic solvents. (D) Phenol partition at different dodecane concentrations. (E) Anisole partition at different dodecane concentrations. (F) Schematic of phenol O-methyltransferase activity by CVOMT1. (G) Phenol consumption in 24 h. (H) Anisole production after 24 h. Abbreviations: (PHE) phenol, (ANI) anisole, (SAM) S-adenosylmethionine, (SAH) S-adenosyl-L-homocysteine, (MET) methionine, (EV) empty vector. Data points and bar graphs represent the mean value ± SD from three independent experiments (*p ≤ 0.05, **p ≤ 0.01, ****p ≤ 0.0001; unpaired Student’s t-test).

To establish this system, we evaluated four organic solvents commonly applied in two-phase whole-cell biocatalysis (dodecane, ethyl acetate, isopropyl myristate and tributyrin),^10,23–25^ aiming to selectively partition anisole into the organic layer while retaining phenol in the aqueous phase for enzymatic conversion.

When aqueous solutions of phenol or anisole were incubated with each solvent and quantified by HPLC, all four solvents extracted over 95% of the anisole (Figure 1B). In contrast, phenol extraction varied substantially across the solvents, exceeding 95% for ethyl acetate and tributyrin, reaching approximately 90% for isopropyl myristate, and remaining at only 15% for dodecane (Figure 1C). This low partition into dodecane preserves substrate availability in the aqueous broth. Titration of the dodecane fraction from 5% to 50% (v/v) during 24-hour incubations maintained phenol extraction at ∼15% (Figure 1D) while increasing anisole recovery from 23.4% to 98% (Figure 1E). Therefore, we adopted a 50% (v/v) dodecane overlay for all subsequent cultivations.

### *In vivo* Phenol Methylation by CVOMT1

With tolerance and product retention addressed, we tested whether CVOMT1 converts phenol to anisole *in vivo*. We expressed the codon-optimized *cvomt* (AF435007.1) gene from a pSEVAb23 vector in *P. putida*, supplemented the cultures with 1 mM phenol under a 50% (v/v) dodecane overlay (Figure 1F) and measured phenol and anisole after 24 h. The CVOMT1-expressing strain consumed 0.049 mM phenol and produced 0.038 mM anisole, whereas the empty-vector control showed negligible background loss (0.002 mM; Figure 1G,H), confirming the predicted *in vivo* activity of CVOMT1. Because SAM is the obligatory methyl donor, we next tested whether cofactor availability limits conversion by supplementing cultures with 1 mM methionine. This supplementation increased phenol consumption to 0.157 mM and anisole production to 0.062 mM (Figure 1G,H), corresponding to 3.2-fold and 1.6-fold increases, respectively, confirming that SAM availability constrained conversion.

### Growth-Coupled Selection of Phenol Methylation Activity

To engineer CVOMT1 toward higher activity on phenol, we implemented a selection system in which cell growth is directly coupled to SAM-dependent methylation.^26^ This design relies on converting methionine into cysteine via the intermediate homocysteine with the aid of a methyltransferase. Exogenously supplied methionine is converted to SAM, which CVOMT1 consumes to methylate phenol, generating anisole and S-adenosylhomocysteine (SAH). A synthetic reverse transsulfuration pathway then converts SAH through homocysteine into cysteine (Figure 2A). In this design, cellular growth is coupled to the *in vivo* methylation rate of CVOMT1 towards phenol: faster methylation generates more SAH, feeding more homocysteine into the reverse transsulfuration pathway to produce cysteine and support faster growth.

**Figure 2.**
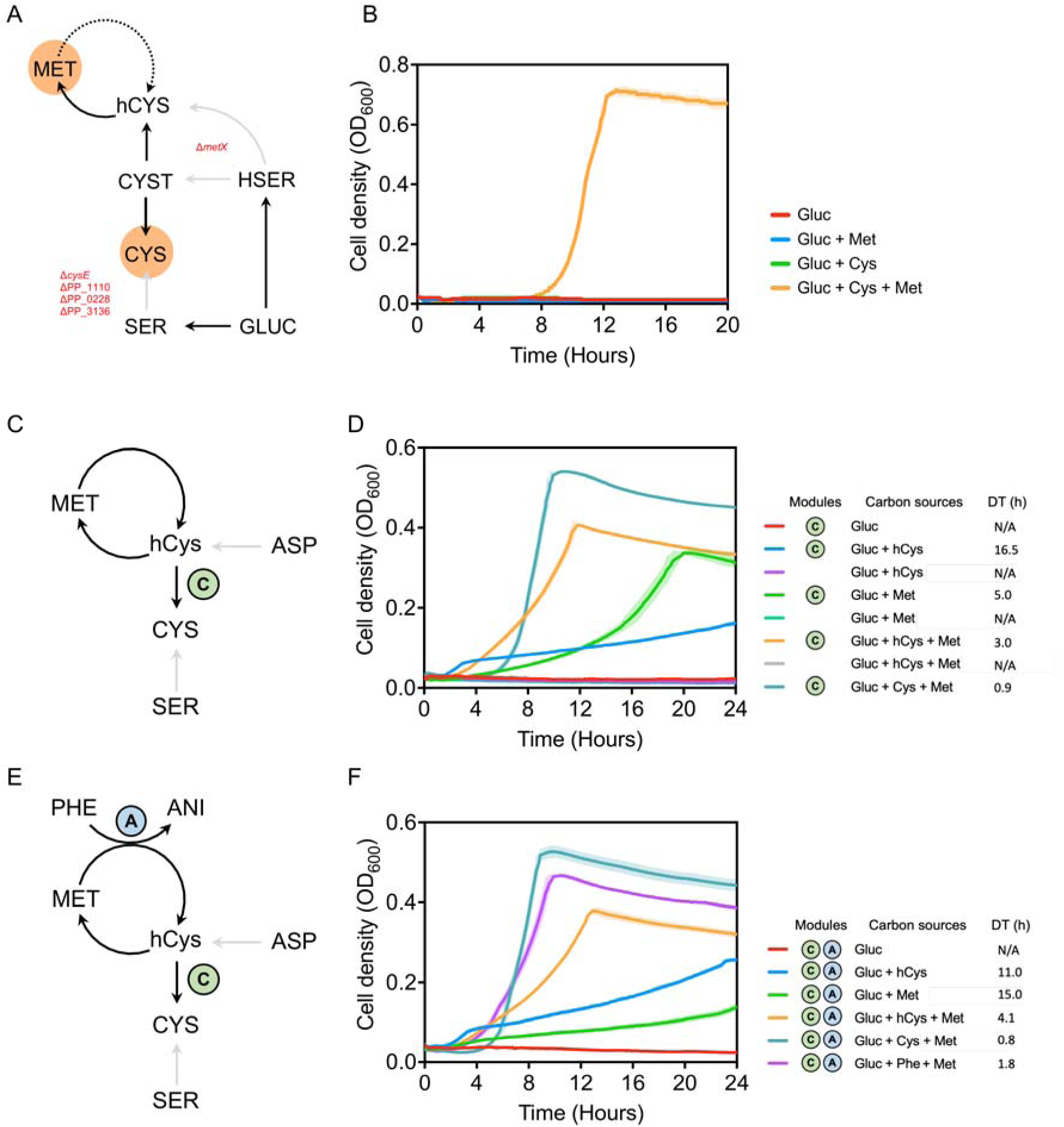
Establishing a growth-coupled methyltransferase selection system. (A) Schematic of the constructed knockouts. (B) Growth curves of CM-Aux; growth occurs only when both cysteine and methionine are supplied. (C) Selection of the C-module in CM-Aux. (D) Growth curves of CM-Aux with the C-module implemented. (E) Selection of the C- and A-modules in CM-Aux. (F) Growth curves of CM-Aux with both the C- and A-modules implemented. Abbreviations: (GLUC) glucose, (CYS) cysteine, (MET) methionine, (hCYS) homocysteine, (CYST) cystathionine, (ASP) aspartate, (SER) serine, (PHE) phenol, (ANI) anisole, (HSER) homoserine. Growth curves represent the mean value ± SD from three independent experiments.

Implementing this platform required a *P. putida* strain auxotrophic for both cysteine and methionine (Figure 2A). Deleting the annotated serine *O*-acetyltransferase genes *cysE* and PP_1110 did not abolish cysteine prototrophy, whereas additionally deleting the paralog PP_0228 did, and PP_3136 was removed as well to tighten the selection (Figure S2A–C). Cysteine can be synthesized not only from serine but also via transsulfuration of cystathionine, an intermediate within methionine biosynthesis. In theory, cysteine could still be produced through this pathway in *P. putida*, yet we observe that the intracellular flux cannot rescue growth (Figure S2D). To completely uncouple methionine and cysteine biosynthesis from the central metabolism, we deleted the *metX* gene, encoding a homoserine *O*-acetyltransferase. The growth of this strain, termed CM-Aux, could only be restored through the addition of both cysteine and methionine, indicating a tight selection system (Figure 2B; details in the Supporting Information, “Construction of the Selection Strain”).

Next, we installed the reverse transsulfuration module (the C-module) to convert homocysteine into cysteine. This two-step pathway combines a cystathionine β-synthase (*cys4* from *Saccharomyces cerevisiae*) and a cystathionine γ-lyase (PA0400 from *Pseudomonas aeruginosa*) cloned into a pSEVAb62 backbone (Figure 2C). Expressing the C-module in CM-Aux rescued growth on homocysteine, yielding a doubling time of 16.5 h on homocysteine alone and 3.0 h upon co-supplementation with methionine (Figure 2D). Growth also occurred with methionine as the sole supplement (doubling time 5.0 h), indicating that native *P. putida* methyltransferases provide sufficient basal SAH flux to restore growth. Although this background activity reduces selection stringency, cells expressing kinetically superior CVOMT1 variants should outgrow slower enzymes. To validate this hypothesis, we introduced the A-module (expressing wild-type *cvomt1*) alongside the C-module in CM-Aux (Figure 2E). In this strain, growth on methionine alone was reduced to a doubling time of 15 h, whereas adding both phenol and methionine restored rapid proliferation with a doubling time of 1.8 h (Figure 2F). This distinct growth acceleration demonstrated that CVOMT1 is catalytically active on phenol *in vivo* and that its activity is successfully coupled to host proliferation.

### Directed Evolution of CVOMT1

Prior to directed evolution of CVOMT1, we calibrated the selection stringency to ensure that improvements in catalytic efficiency could be clearly resolved. Under saturating substrate concentrations (1 mM phenol and 1 mM methionine), wild-type CVOMT1 already supported rapid growth (doubling time 1.8 h; Figure 2F), which would mask differences among higher-performing variants. We therefore titrated both methionine and phenol across the selection strain (Figure 3A,B). Methionine concentrations had a negligible effect, consistent with the low K of CVOMT1 for SAM (5 µM).^17^ In contrast, growth was strongly sensitive to phenol across the sub-millimolar range, identifying the non-native substrate phenol as the growth-limiting factor. We therefore selected 0.1 mM phenol as the screening condition, ensuring that small gains in catalytic turnover would translate into measurable growth advantages.

**Figure 3.**
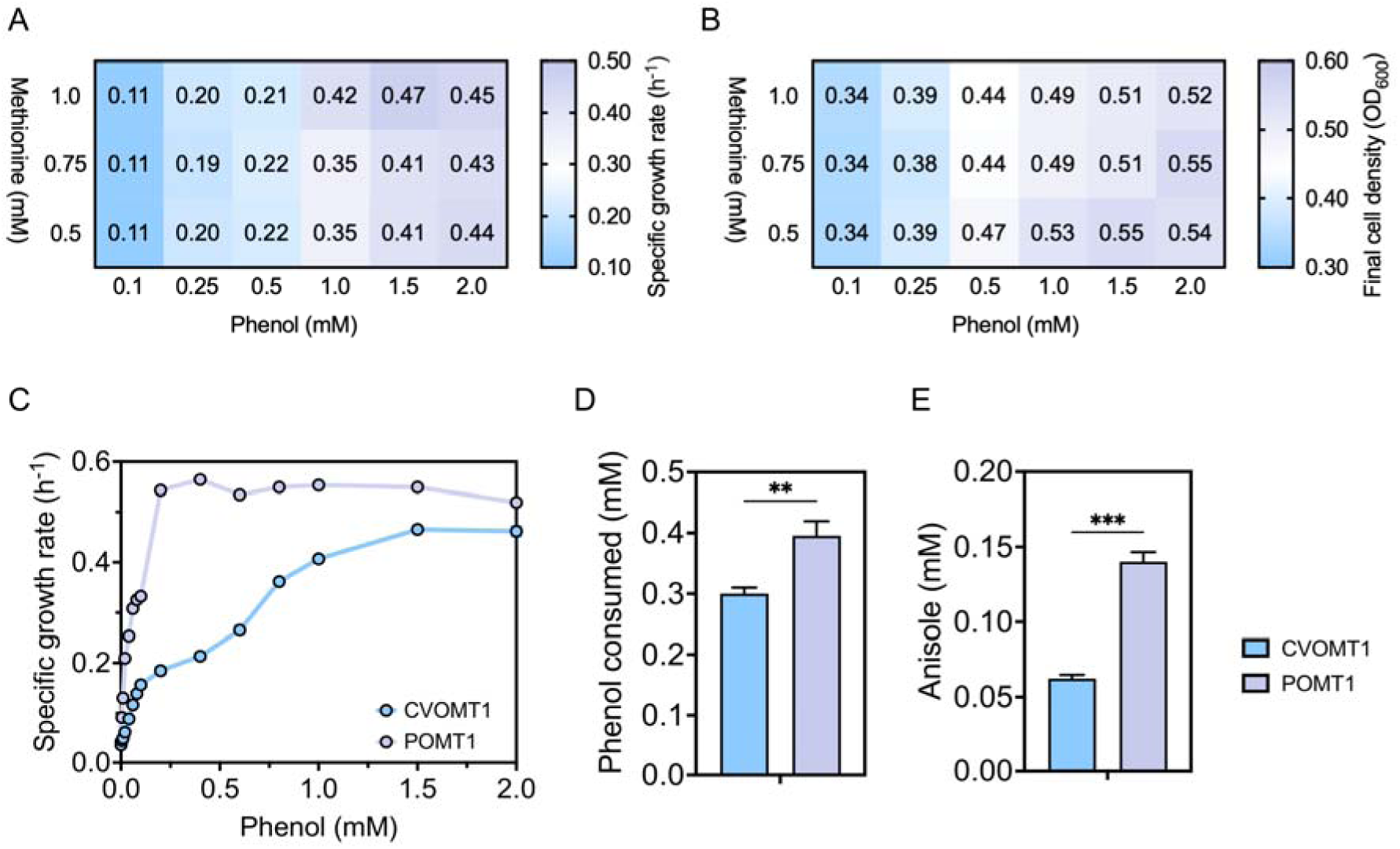
Establishment and characterization of POMT1. (A) Heatmap of specific growth rates of CM-Aux expressing the C- and A-modules at different methionine and phenol concentrations. (B) Heatmap of final cell densities under the same conditions. (C) Specific growth rates of CM-Aux equipped with CVOMT1 or POMT1 across phenol concentrations. (D) Phenol consumption, CVOMT1 vs. POMT1. (E) Anisole production, CVOMT1 vs. POMT1. Data points represent the mean value ± SD from three independent experiments (n = 3; **p ≤ 0.01, ***p ≤ 0.001; unpaired two-tailed Student’s t-test). See also Figure S3 (selection stringency), Figure S4 and Tables S5 and S6 (Boltz-2 predictions).

To optimize CVOMT1, we considered both random mutagenesis and adaptive laboratory evolution (ALE). Although ALE is an efficient tool for strain improvement, it carries the risk of background rescue, where native host methyltransferases evolve to bypass the selection system.^26^ Therefore, to obtain improved variants, we generated a *cvomt1* library by error-prone PCR and introduced it into CM-Aux. Following library’s enrichment in liquid minimal medium (10 mM glucose, 1 mM methionine, 0.1 mM phenol), we plated the culture onto matching selective agar and picked 92 of the largest colonies for individual characterization in microtiter plates. At 0.1 mM phenol, every selected variant grew faster than the wild-type control (Figure S3A), but the isolates could not be resolved from one another. In contrast, reducing the phenol concentration to 0.01 mM separated them into a broad spectrum of distinct growth phenotypes (Figure S3B), demonstrating that selection stringency is a tunable parameter of this platform. Whole-plasmid nanopore sequencing of the five fastest-growing isolates revealed that all five carried the same substitution, aspartic acid 264 to asparagine (D264N), with no other mutations in *cvomt1*. We designated this variant POMT1 (phenol *O*-methyltransferase 1).

Then, we compared POMT1 with wild-type CVOMT1, first in the selection strain and then in production strains. In CM-Aux, POMT1 consistently outgrew wild-type CVOMT1 across a range of phenol concentrations and reached its maximum growth rate at 0.2 mM phenol (Figure 3C). To test whether this advantage translates into product titer, we transferred both enzymes into production strains of *P. putida* and supplied 1 mM phenol under a 50% dodecane overlay for 24 h. POMT1 consumed 0.39 mM phenol, 30% more than the 0.30 mM consumed by CVOMT1 (Figure 3D), and produced 0.14 mM anisole, a 2.3-fold increase over the 0.06 mM obtained with the wild-type enzyme (Figure 3E), so a single round of selection delivered a substantially improved biocatalyst. The gap between phenol consumed and anisole recovered likely reflects residual volatilization and extraction losses.

To investigate the structural basis of this enhanced activity, we examined the active-site context of position 264. Residue 264 lies adjacent to His-263, the putative catalytic base responsible for deprotonating the substrate hydroxyl group.^27,28^ To explore this mechanism, we generated co-complex models of CVOMT1 and POMT1 with SAM and phenol using Boltz-2, a structure and binding-affinity prediction model.^29^ While predicted binding affinities were indistinguishable between the variants (Table S5), the sampled substrate poses differed markedly. Across 50 models per enzyme, the phenol oxygen lay within 3.5 Å of the SAM methyl group in 49 POMT1 models compared to 30 in CVOMT1, with the side chain of residue 264 positioned within 3.4 Å of the phenol oxygen in 47 and 17 models, respectively, although the top-ranked model of each phenol run was reactive in both enzymes (Table S6, Figure S4). We therefore hypothesize that an amide, unlike a carboxylate, can hydrogen-bond the phenol hydroxyl without competing with His-263 for the proton or disfavoring the developing phenolate. Asn occurs at this position in 12.6% of 3,597 aligned homologs (Table S7), and single residues can determine substrate preference in plant *O*-methyltransferases.^17,30^ However, this hypothesis rests on computational predictions and mechanistic reasoning. Because co-folding models do not reliably capture the subtle effects of point mutations,^31^ and whole-cell assays cannot separate higher catalytic turnover from increased enzyme abundance, future validation with purified enzyme will be required.

To test whether natural enzymes could match POMT1, we evaluated a panel of five candidate *O*-methyltransferases identified from literature reports, sequence mining with EnzymeMiner, and structural search using Foldseek.^32–35^ When evaluated in the CM-Aux selection strain, only POMT1 sustained rapid growth at 0.1 mM phenol, which represents the low-concentration regime relevant to *de novo* pathway integration (Figure S5). Notably, the *Geodermatophilus obscurus* enzyme, which natively carries the D264N-equivalent Asn residue, showed negligible activity, indicating that this residue alone is insufficient (Supporting Information).

### Comparison of the Salicylate and 4-Hydroxybenzoate Routes to Phenol

Achieving *de novo* biosynthesis of anisole requires establishing an efficient metabolic route from glucose to the precursor phenol. From the central shikimate intermediate chorismate, phenol can be synthesized via non-oxidative decarboxylation of either 4-hydroxybenzoate (4HB) or salicylate (SAL) (Figure 4A). Although phenol can also be produced via tyrosine, genome-scale metabolic simulations using the *P. putida* genome-scale model iJN1462^36^ predicted comparable theoretical yields across the pathways (0.534 mol anisole per mol glucose for tyrosine versus 0.531 for 4HB and SAL). We favored the 4HB and SAL routes because the tyrosine pathway requires more enzymatic steps, involves intricate allosteric regulation,^37^ and suffers from severe feedback inhibition of tyrosine phenol-lyase by phenol.^18^ We therefore evaluated both candidate routes in *P. putida*.

**Figure 4.**
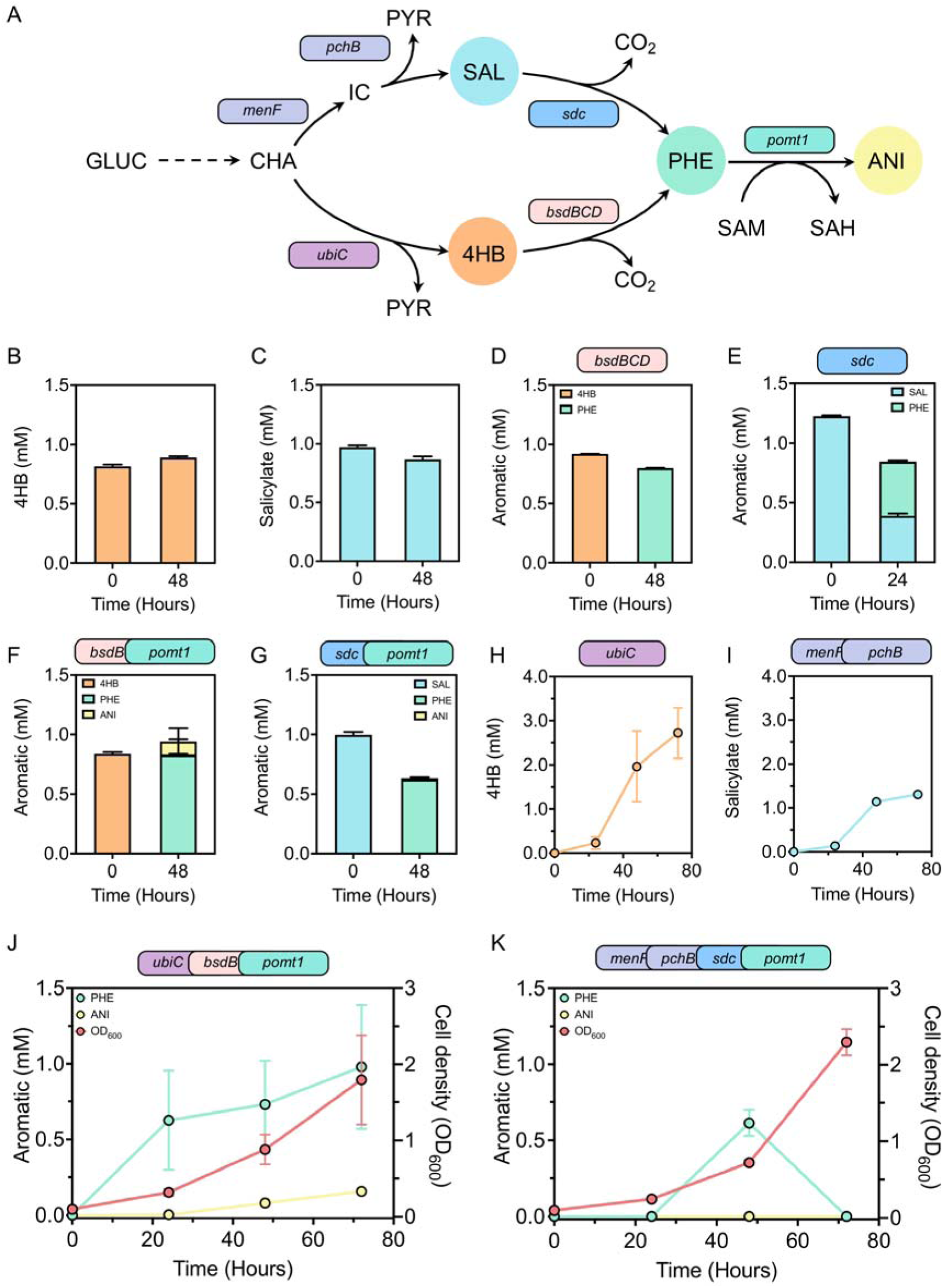
Precursor route selection and *de novo* production of anisole. (A) Schematic of the anisole production pathway in *P. putida* KT2440; chemicals and their corresponding overexpressed genes share color coding. (B) 4HB degradation by *P. putida* Δ*pobA*. (C) SAL degradation by *P. putida* Δ*pobA*. (D) Phenol production from 4HB. (E) Phenol production from SAL. (F) Anisole production from 4HB. (G) Anisole production from SAL. (H) 4HB production from glucose. (I) SAL production from glucose. (J) Anisole production from glucose via the 4HB pathway. (K) Anisole production from glucose via the SAL pathway. Abbreviations: (GLUC) glucose, (CHA) chorismate, (PYR) pyruvate, (IC) isochorismate, (4HB) 4-hydroxybenzoate, (SAL) salicylate, (PHE) phenol, (ANI) anisole, (SAM) S-adenosylmethionine, (SAH) S-adenosyl-L-homocysteine, (CO_2_) carbon dioxide. Genes: *ubiC* (chorismate pyruvate lyase), *menF* (isochorismate synthase), *pchB* (isochorismate pyruvate lyase), *sdc* (salicylate decarboxylase), *bsdBCD* (4-hydroxybenzoate decarboxylase), *pomt1* (phenol *O*-methyltransferase). Data points represent the mean value ± SD from three independent experiments.

To compare the routes, we first stabilized the precursors and then expressed the decarboxylases. We deleted *pobA* (encoding p-hydroxybenzoate hydroxylase) to prevent 4HB catabolism, whereas *P. putida* lacks native SAL degradation, and observed that neither precursor was degraded over the course of two days (Figure 4B,C). For the 4HB route, we attempted to express the *Bacillus subtilis bsdBCD* operon^38^ (encoding a multi-subunit, non-oxidative decarboxylase complex responsible for decarboxylating 4HB into phenol) from a high-copy plasmid, yet this caused severe growth arrest, likely from acute phenol accumulation. Therefore, we integrated *bsdBCD* into the chromosome through a Cre/lox landing pad under T7 RNA polymerase control, which gave complete conversion of 4HB to phenol over 48 h (Figure 4D). In the SAL route, plasmid-borne *Trichosporon moniliiforme* salicylate decarboxylase (SDC)^39^ acted rapidly (Figure 4E) but recovered carbon incompletely: only 37.2% of the SAL was converted to phenol after 24 h, 31.6% remained unreacted and total aromatic recovery fell by 31.2%, which suggests off-target turnover of intermediates by SDC (Figure 4C).

We then compared the routes by anisole formation, co-expressing *pomt1* with each decarboxylase from a pSEVAb23 plasmid. In the 4HB route, POMT1 alleviated the plasmid-associated toxicity of *bsdBCD* alone, consistent with POMT1 lowering intracellular phenol by converting it to anisole. With 1 mM 4HB, this strain produced 0.825 mM phenol and 0.110 mM anisole after 48 h (Figure 4F), and the total aromatic pool exceeded the supplied 4HB by 11.75%, consistent with endogenous 4HB influx from ubiquinone biosynthesis in the Δ*pobA* background (Figure 4B). In contrast, the SAL route converted the supplied SAL completely to phenol (0.62 mM; Figure 4G) but gave no anisole and an incomplete mass balance, which suggests that SDC may convert phenol and divert intermediates before methyl transfer by POMT1.

Finally, we tested both routes in *de novo* production from glucose, starting from a background engineered for higher shikimate-pathway flux (characterized below). We expressed a feedback-resistant chorismate pyruvate lyase (*ubiC*^E31Q/M34V^ from *E. coli*)^40^ for the 4HB route or a synthetic *menF*–*pchB* operon (*E. coli menF*, *P. aeruginosa pchB*) for the SAL route, both under arabinose-inducible control. From 50 mM glucose, the strains produced 2.72 mM 4HB and 1.31 mM SAL after 72 h (Figure 4H,I). With the evolved *pomt1* added, the 4HB route made 0.62 mM phenol within 24 h and 0.98 mM by 72 h while anisole accumulated steadily to 0.16 mM (Figure 4J), whereas the SAL route accumulated phenol only transiently (0.61 mM at 48 h) and gave no detectable anisole (Figure 4K), the same signature of phenol consumption by SDC. We therefore selected the 4HB route for all subsequent *de novo* work, because it accumulated phenol stably and yielded detectable anisole.

### Expression, Cofactor Supply, and Precursor Engineering for *De Novo* Anisole Production

In the glucose-fed 4HB strain, phenol accumulated to 0.98 mM against only 0.16 mM anisole, so the catalytic turnover of POMT1 still constrained throughput. Because CVOMT1 originates from the plant *O. basilicum*, we first tested whether N-terminal affinity tags could improve its folding in *P. putida*.^41^ Both a 6×His-tag and a FLAG-tag increased phenol consumption (1.61-fold and 2.93-fold, respectively) and anisole production (2.62-fold and 4.28-fold) relative to untagged POMT1 (Figure S6A,B), which suggests that folding or solubility limited the expression of the untagged enzyme, and we used FLAG-POMT1 in all subsequent strains. Comparing constitutive promoters and plasmid backbones, we found that promoter identity made negligible difference, whereas the higher-copy pSEVAb24 backbone^42^ lowered phenol consumption and anisole production by 2.05-fold and 1.89-fold, respectively (Figure S6C), so gene dosage must be balanced rather than maximized.

We next examined cofactor supply as another possible bottleneck. Co-expressing the SAM synthase gene *metK* across 0 to 2 mM methionine did not raise the final anisole titer (Figures S6E and S7D), unlike the benefit of methionine seen with unevolved CVOMT1 (Figure 1G,H), and higher methionine slowed phenol consumption in *metK*-overexpressing strains (1.82 mM without methionine versus 1.29 mM at 2 mM, from 2 mM phenol within 24 h; Figure S7B), consistent with feedback inhibition rather than substrate limitation. Nevertheless, all strains fully consumed the supplied phenol within 48 h (Figure S7). To test whether SAH recycling limits conversion, we also introduced the orthogonal *E. coli mtn*–*luxS* module, which recycles SAH to homocysteine, and again observed no improvement (Figure S6F). These matched negative controls indicate that neither SAM synthesis nor SAH recycling limits conversion, which narrows optimization to the catalytic turnover of POMT1 (Supporting Information).

Because SAM supply and SAH recycling did not appear to limit throughput, we next increased the precursor supply through the shikimate pathway. We first raised the supply of phosphoenolpyruvate (PEP) by deleting the competing PEP-consuming enzymes (Δ*pyk* Δ*pykA* Δ*ppc*), together with Δ*pobA* and Δ*quiC* to block competing aromatic catabolic branches (Figure 5A). In 50 mL tubes with an aerated cap and a 1:1 (v/v) dodecane overlay, this strain produced 0.23 mM anisole, a 64.3% increase over its control parent (0.14 mM; Figure 5B), consistent with enriched intracellular PEP promoting pathway throughput. Because the shikimate pathway also requires erythrose-4-phosphate (E4P), which is scarce in *P. putida*,^43^ we then integrated *E. coli* transketolase (*tktA*) and transaldolase (*talB*) at the chromosomal *att*Tn7 locus.^44,45^ However, this intervention had a negligible effect on anisole production (0.23 mM) and only slightly increased total aromatics through accumulation of unconverted phenol (∼0.06 mM; total aromatics 0.28 mM; Figure 5B).

**Figure 5.**
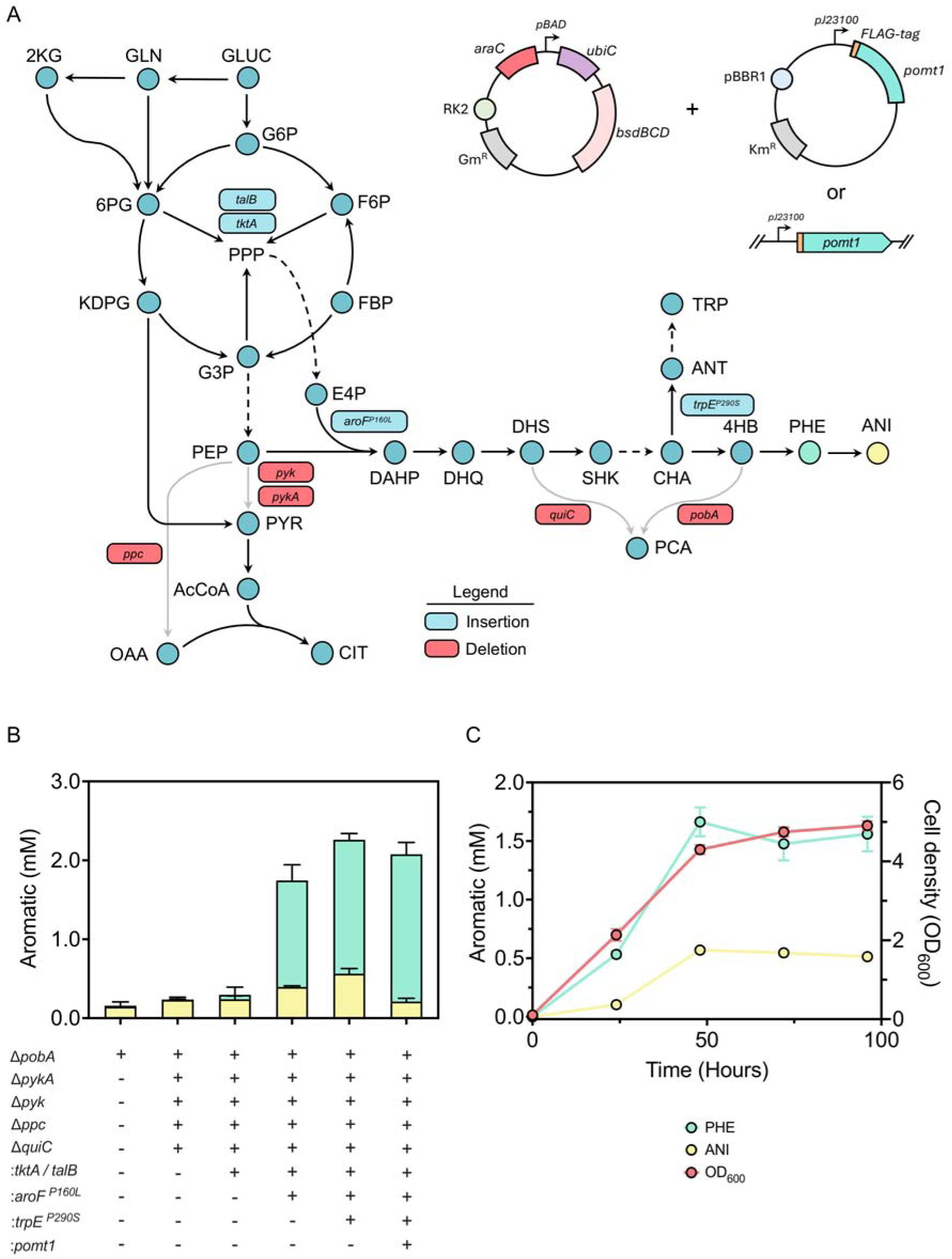
Optimization of precursor biosynthesis for anisole production. (A) Schematic of the anisole production pathway in *P. putida* KT2440; gray arrows marked with a red box denote genetic deletions. (B) Anisole titers of optimized production strains. (C) *De novo* anisole biosynthesis over four days. Abbreviations: (GLUC) glucose, (GLN) gluconate, (2KG) 2-ketogluconate, (G6P) glucose-6-phosphate, (F6P) fructose-6-phosphate, (FBP) fructose bisphosphate, (6PG) 6-phosphogluconate, (KDPG) 2-keto-3-deoxy-6-phosphogluconate, (G3P) glyceraldehyde-3-phosphate, (PEP) phosphoenolpyruvate, (E4P) erythrose-4-phosphate, (PYR) pyruvate, (AcCoA) acetyl-coenzyme A, (OAA) oxaloacetate, (CIT) citrate, (DAHP) 3-deoxy-D-arabino-heptulosonate-7-phosphate, (DHQ) 3-dehydroquinate, (DHS) 3-dehydroshikimate, (SHK) shikimate, (CHA) chorismate, (PCA) protocatechuate, (ANT) anthranilate, (TRP) L-tryptophan, (4HB) 4-hydroxybenzoate, (PHE) phenol, (ANI) anisole. Genes: *pyk*/*pykA* (pyruvate kinase), *ppc* (PEP carboxylase), *quiC* (3-dehydroshikimate dehydratase), *pobA* (*p*-hydroxybenzoate hydroxylase), *talB* (transaldolase), *tktA* (transketolase), *aroF-I^P160L^* (feedback-resistant DAHP synthase), *trpE*^P290S^ (anthranilate synthase component I, reduced-flux variant), *pomt1* (phenol *O*-methyltransferase). Data points represent the mean value ± SD from three independent experiments.

Next, we targeted the entry point of the shikimate pathway, where relieving feedback inhibition is a well-established strategy for increasing the aromatic flux.^46^ Following a feedback-resistant 3-deoxy-D-arabino-heptulosonate-7-phosphate (DAHP) synthase variant from *Pseudomonas taiwanensis* (AroF-I P148L),^47^ we introduced the homologous P160L substitution (adjusted for 12 additional N-terminal residues) into the *P. putida* enzyme, which lifted titers to 0.39 mM anisole and 1.36 mM phenol, the largest single gain. Adding *trpE* P290S, which redirects chorismate away from tryptophan biosynthesis toward 4HB,^48^ raised them further to 0.56 mM anisole and 1.7 mM phenol. Because plasmid-based expression requires antibiotic selection, which industrial processes prefer to avoid,^49^ we also tested a chromosomal copy of POMT1. Integrating FLAG-POMT1 under the constitutive J23100 promoter downstream of PP_5322, a locus with high basal expression and low fitness cost,^50^ left phenol accumulation unaffected but lowered anisole from 0.56 to 0.20 mM.

Finally, we combined plasmid-expressed FLAG-POMT1 with all upstream modifications (Figure 5B) and cultivated the resulting strain in shake flasks on 50 mM glucose for four days (Figure 5C). Anisole accumulation peaked at 0.57 mM after 48 h alongside 1.66 mM residual phenol. Together with the cofactor diagnostics that excluded SAM limitation, these results identify the turnover of the terminal methyltransferase as the primary remaining bottleneck.

Although this establishes proof-of-concept for the *de novo* biosynthesis of anisole from glucose in *P. putida*, current titers (62 mg L ¹) remain below levels required to compete with petroleum-derived chemical synthesis. Upstream metabolic engineering successfully decoupled precursor supply from product formation, yet the persistent accumulation of unconverted phenol (1.66 mM), combined with cofactor diagnostics that excluded SAM limitation, demonstrates that the terminal methylation step remains the primary kinetic bottleneck. Future iterations will focus on further directed evolution of POMT1 or identifying orthogonal methyltransferases with higher catalytic turnover, alongside multi-copy chromosomal integration to achieve stable, antibiotic-free expression. Table S8 summarizes the stepwise progression of phenol and anisole titers from the first proof-of-concept conversion to the complete de novo production strain.

## EXPERIMENTAL SECTION

### Strains, Plasmids, and Culture Conditions

Strains, plasmids, primers and synthetic genes are listed in Tables S1–S3. *P. putida* and *E. coli* were grown at 30 and 37 °C, respectively. Growth and production experiments used M9 minimal medium with trace elements, inoculated at OD_600_ 0.1 from washed LB precultures, with 10 mM glucose in plate-reader experiments and 50 mM glucose in 250 mL flasks (25 mL working volume, 200 rpm, 30 °C). Unless stated otherwise, cysteine, methionine, homocysteine, phenol and 4HB were added at 1 mM, and 2 mM arabinose was added at inoculation where genes were arabinose-inducible (*ubiC*, *menF*–*pchB, sdc, bsdBCD*). Medium compositions are given in the Supporting Information.

### Plasmid and Strain Construction

Plasmids were assembled by SevaBrick assembly^42^ with genes codon-optimized using JCat,^51^ and genomic edits followed the I-SceI-based protocol of Wirth *et al.*^52^ Constructs were verified by Sanger sequencing and, for the five fastest-growing isolates, by whole-plasmid nanopore sequencing (Supporting Information).

### Tolerance, Evaporation, and Two-Phase Assays

Growth of *P. putida* with 0–5 mM phenol or anisole was followed in the plate reader (Figure S1), and abiotic anisole loss from aqueous solution was measured by HPLC after 16 h (Figure 1A). Solvent partitioning used 1 mM aqueous phenol or anisole mixed 1:1 with each solvent (1 h, 600 rpm, 30 °C), and the dodecane series (0–100% v/v) used a 25 mL total volume (24 h, 30 °C, 250 rpm). Bioconversions (Figures 1G,H, 3D,E, 4F, S6 and S7) were run in 250 mL flasks with a 25 mL working volume containing 50% (v/v) dodecane and substrate (1 mM phenol or 4HB unless stated) added at t = 0, and both phases were analyzed by HPLC. Reactor-tube assays used 50 mL cell reactor tubes with an aerated cap containing 5 mL medium and 5 mL dodecane.

### Analytical Methods

Phenol, anisole, 4-hydroxybenzoate and salicylate were quantified by HPLC on a C18 column with UV detection at 271 nm and identified by retention time against authentic standards, whereas glucose was quantified by HPLC with refractive-index detection (details in the Supporting Information).

### Growth-Coupled Selection and Screening

A *cvomt1* library generated by error-prone PCR was cloned into pSEVAb23, introduced into CM-Aux carrying the C-module, enriched in M9 with 10 mM glucose, 1 mM methionine and 0.1 mM phenol, and plated. Ninety-two large colonies were screened in microtiter plates at 0.1 and 0.01 mM phenol (details in the Supporting Information).

### Computational Analyses

Flux balance analysis used the genome-scale model iJN1462^36^ extended with the heterologous reactions. Co-complexes of CVOMT1 and POMT1 with SAM and phenol or chavicol were predicted with Boltz-2 (v2.2.1)^29^ from one shared sequence alignment (two runs of 25 models each). A model was called reactive when the SAM methyl carbon lay within 3.5 Å of the substrate oxygen.

### Statistical Analysis

All reported experiments are independent biological triplicates unless otherwise noted. Values are means ± standard deviation, and significance was assessed with an unpaired Student’s *t*-test.

## ASSOCIATED CONTENT

### Data Availability Statement

The data supporting this article, including strain and plasmid sequences and Boltz-2 prediction summaries, are available in the article and its Supporting Information.

### Supporting Information

The Supporting Information is available free of charge at https://pubs.acs.org/doi/10.1021/acssuschemeng.XXXXXXX. Supplementary methods and results, Figures S1–S7 and Table S8 (PDF); Tables S1–S7: strains, plasmids, primers, gene sequences and computational outputs (XLSX).

## Author Contributions

L.B. and C.B. conceived and supervised the study and contributed equally to this work. L.B., D.R.G., C.L.R., J.E.E.M and C.B. designed and performed the experiments and analyzed the data. V.A.P.M.d.S. contributed to supervision, funding acquisition, and manuscript revision. L.B. and C.B. wrote the manuscript with input from all authors. All authors have given approval to the final version of the manuscript.

## Funding

We acknowledge the financial support by the Europe Horizon 2020 projects IBISBA (grant numbers 730976 and 871118) and the Horizon Europe projects BIOS (grant number 101070281) and Bioindustry 4.0 (grant number 101094287).

## Notes

L.B., C.B. and V.A.P.M.d.S. are inventors on patent application WO 2024/253523 (“Biological anisole production”), filed by Wageningen University & Research. The other authors declare no competing financial interest.

## ACKNOWLEDGMENTS

We thank Dr. Maria Martin Pascual for kindly providing the pGNW-aroF-I(P160L) plasmid.

