## Supporting information for "Establishing *de novo* microbial biosynthesis of anisole"

**Contents**

**Supplementary Methods** – culture conditions, construction, assays, analytics, screening, flux balance analysis, structure prediction and statistics

**Supplementary Results** – construction of the selection strain; benchmarking of natural *O*-methyltransferases; N-terminal tags, expression context and cofactor supply; stepwise progression of titers

**Figures S1–S7**

**Table S8** (this document); **Tables S1–S7** are provided in the accompanying Excel file: S1 strains and plasmids, S2 primers, S3 codon-optimized genes, S4 Selenzyme output, S5 Boltz-2 affinity-head output, S6 Boltz-2 pose occupancy, S7 tolerance of Asn at position 264

**References**

### SUPPLEMENTARY METHODS

#### Media, Antibiotics, and Culture Conditions

All strains and plasmids used in this study are listed in Table S1; primers used for plasmid construction and gene deletions are listed in Table S2. *P. putida* and *E. coli* cultures were incubated at 30 °C and 37 °C, respectively. For cloning, both species were propagated in Lysogeny Broth (LB; 10 g $L^{-1}$ NaCl, 10 g $L^{-1}$ tryptone, 5 g $L^{-1}$ yeast extract), solidified with 1.5% (w/v) agar where required. Antibiotics were used at kanamycin (Km) 50 µg $mL^{-1}$, gentamycin (Gm) 10 µg $mL^{-1}$, chloramphenicol (Cm) 30 µg $mL^{-1}$ and apramycin (Apra) 50 µg $mL^{-1}$. Growth and production experiments used M9 minimal medium (per liter: 3.88 g $K_2HPO_4$, 1.63 g $NaH_2PO_4$, 2.0 g $(NH_4)_2SO_4$, pH 7.0) supplemented with trace elements (per liter: 10 mg EDTA, 0.1 g $MgCl_2·6H_2O$, 2 mg $ZnSO_4·7H_2O$, 1 mg $CaCl_2·2H_2O$, 5 mg $FeSO_4·7H_2O$, 0.2 mg $Na_2MoO_4·2H_2O$, 0.2 mg $CuSO_4·5H_2O$, 0.4 mg $CoCl_2·6H_2O$, 1 mg $MnCl_2·2H_2O$). Strains were precultured in 10 mL LB with the appropriate antibiotics, washed twice in carbon-source-free M9 medium, and diluted to $OD_{600}$ 0.1 to start each experiment. Plate-reader experiments used 10 mM glucose; flask experiments used 250 mL Erlenmeyer flasks with a 25 mL working volume and 50 mM glucose. Unless stated otherwise, supplementary carbon sources (cysteine, methionine, homocysteine, phenol, SAL, 4HB) were added at 1 mM. Plate-reader experiments were run in 200 µL M9 medium on an ELx808 plate reader (BioTek) with continuous shaking, recording $OD_{600}$ every three minutes. Flask cultures were incubated at 200 rpm and 30 °C. Where pathway genes were under arabinose-inducible control (*ubiC*, *menF–pchB*), 2 mM arabinose was added at inoculation.

#### Plasmid and Strain Construction

Plasmids were assembled using SevaBrick Assembly.[1] DNA fragments were amplified with Q5® Hot Start High-Fidelity DNA Polymerase (New England Biolabs). Genes were codon-optimized with JCat[2] and synthesized by GenScript (Table S3). Plasmids were transformed by heat shock into chemically competent *E. coli* DH5α λpir, selected on LB agar with the appropriate antibiotic, screened by colony PCR (Phire Hot Start II DNA Polymerase, Thermo Fisher Scientific), and verified by Sanger sequencing (Macrogen) before electroporation into *P. putida*. Genomic deletions and insertions followed the I-SceI-based protocol of Wirth *et al.*:[3] ~500 bp homology arms flanking each target gene were cloned into the non-replicative pGNW vector, propagated in *E. coli* DH5α λpir, and transformed into *P. putida* by electroporation, with co-integrants selected on LB + Km. Co-integrated strains were then transformed with the helper plasmid pQURE6-H and plated on LB + Gm with 2 mM 3-methylbenzoic acid to induce the XylS-dependent Pm promoter driving the I-SceI homing nuclease, resolving the co-integrate. Successful deletions were confirmed by PCR and Sanger sequencing, after which pQURE6-H was cured by removing selective pressure and confirmed lost by gentamycin sensitivity. Plasmids from the five fastest-growing isolates of the growth-coupled selection were recovered and verified by whole-plasmid nanopore sequencing.

#### Tolerance, Evaporation, Two-Phase Bioconversion, and Reactor-Tube Assays

Growth of *P. putida* KT2440 with phenol or anisole (0, 0.25, 0.5, 1, 2, 3, 4 and 5 mM) was followed in the plate reader under the conditions described above, and specific growth rates were calculated from the $OD_{600}$ curves (Figure S1). Abiotic loss of anisole from aqueous solution (Figure 1A) was determined by incubating 25 mL of 2 mM anisole in M9 minimal medium in 250 mL Erlenmeyer flasks at 30 °C and 200 rpm [or 250 rpm], with samples taken at t = 0 and 16 h for HPLC quantification. Two-phase bioconversions (Figures 1G,H, 3D,E and 4F, S6 and S7) were run in 250 mL Erlenmeyer flasks with a 25 mL working volume containing 50% (v/v) dodecane, with substrate (1 mM phenol, SAL or 4HB unless stated otherwise) added at t = 0 and both phases sampled for HPLC analysis. Reactor-tube assays (Figure 5B) used aerated-cap 50 mL tubes with 5 mL medium and 5 mL dodecane.

**Organic-Phase Extraction and Partition Ratio Determination**

Phenol (Sigma-Aldrich) and anisole (Thermo Scientific) were diluted in water to 1 mM. 900 µL of each dilution was combined with an equal volume of the test solvent (dodecane [VWR], ethyl acetate [Sigma-Aldrich], isopropyl myristate or tributyrin [Thermo Scientific]) in 2 mL Eppendorf tubes, incubated for 1 hour at 600 rpm and 30 °C, and centrifuged for 5 minutes at maximum speed; 200 µL from each phase was analyzed by HPLC. For dodecane ratio optimization, six fractions (0%, 5%, 15%, 30%, 50% and 100% v/v) were tested using Milli-Q water as the aqueous phase and 1 mM phenol and anisole (based on a 25 mL total working volume). An initial 1 mL aqueous-phase sample was taken at t = 0 before dodecane addition; flasks were then incubated at 30 °C and 250 rpm for 24 hours, after which the phases were separated by centrifugation (4000 rpm, 15 min) and sampled for HPLC analysis of both layers.

**Analytical Methods**

Cell growth was monitored by $OD_{600}$ using an OD600 DiluPhotometer (IMPLEN) or a ELx808 plate reader (BioTek) Glucose was quantified by HPLC (Thermo Fisher Scientific) with an Aminex HPX-87H column, 5 mM $H_2SO_4$ mobile phase at 0.6 mL $min^{-1}$, 60 °C column temperature, and refractive-index detection (Shodex RI-101). 4-Hydroxybenzoate, salicylate, phenol and anisole were quantified by HPLC (Shimadzu) on a C18 column (4.6 mm × 250 mm) with UV/vis detection at 271 nm, using a gradient of Milli-Q water (A), 100 mM formic acid (B) and acetonitrile (C) at 1 mL $min^{-1}$ and 30 °C (0–5 min: 55% A, 10% B, 35% C; ramped to 10% A, 10% B, 80% C by 10 min and held to 15 min; returned to initial conditions by 16 min; total run time 18 min). Peaks were identified by comparing retention times with those of authentic standards, and calibration curves used 99% purity standards (Sigma-Aldrich).

**High-Throughput Screening of CVOMT1 Variants**

A *cvomt1* mutant library was generated by error-prone PCR using the GeneMorph II Random Mutagenesis Kit (Agilent Technologies) from 50 ng template DNA with primers CVOMT_ALE_FW/RV over 30 thermal cycles, gel-purified, and cloned into pSEVAb23. The library was electroporated into CM-Aux carrying the C-module, grown overnight in LB with antibiotics, washed twice in carbon-source-free M9, and reinoculated at $OD_{600}$ 0.1 in M9 with 10 mM glucose, 1 mM methionine and 0.1 mM phenol for overnight enrichment. The enriched culture was plated on matching M9 agar; after 2 days, 92 large colonies were picked into 96-well plates (200 µL LB with antibiotics; two wells of unevolved CVOMT1 as positive control, two blank wells as negative control) and grown for 24 hours at 30 °C (ELx808 plate reader). 5 µL of each culture was then transferred to fresh M9 (10 mM glucose, 1 mM methionine, 0.1 mM phenol) and monitored by $OD_{600}$ every three minutes over 24 hours at 30 °C. The isolates were also characterized at 0.01 mM phenol (Figure S3B).

#### Genome-Scale Flux Balance Analysis

The genome-scale metabolic model iJN1462 of *P. putida* was extended with reactions for phenol O-methyltransferase, salicylate decarboxylase, 4-hydroxybenzoate decarboxylase, and tyrosine phenol-lyase. Simulations were performed under aerobic conditions with the glucose uptake rate constrained to 10 mmol $gCDW^{-1}$ $h^{-1}$. Flux balance analysis was used to calculate the maximum theoretical anisole yield per route by setting anisole export as the objective function, resulting in 0.534 mol $mol^{-1}$ glucose for the tyrosine route and 0.531 mol $mol^{-1}$ glucose for the 4HB and SAL routes.

#### Structure Prediction and Sequence Analysis

Co-complex structures of CVOMT1 and POMT1 with SAM and either phenol or chavicol were predicted using Boltz-2 (version 2.2.1)[4] on the Tamarind platform. SAM was supplied by its CCD code and the substrate as a SMILES string (phenol: Oc1ccccc1; chavicol: C=CCc1ccc(O)cc1), with the substrate as the binder chain for the affinity head. Both enzymes were run with one supplied sequence alignment, so that they differ only at residue 264. The alignment was built by the Tamarind Boltz-2 service for the CVOMT1 sequence with MMseqs2 searches against UniRef30 and the ColabFold environmental database.[5,6] It holds the query and 12,625 homologous sequences (12,627 records, because the query heads each of the two database blocks). For POMT1, the query residue at position 264 was changed to Asn and the file was otherwise identical. Each complex was predicted in two independent runs (seeds 2 and 3) of 25 diffusion samples with three recycling steps and potentials enabled. The affinity head returns one log10 IC50 (μM) value per run, which was converted to kcal/mol equivalents as (log10 IC50 − 6) × 1.364. These values are not binding free energies. Inter-atomic distances were measured in each of the 50 models per system. A model was called reactive when the SAM methyl carbon lay within 3.5 Å of the substrate oxygen. Contact with residue 264 was defined as a side-chain oxygen or nitrogen within 3.4 Å of the substrate oxygen, and contact with His-263 as its Nε2 atom within 3.2 Å. The residue aligned to position 264 was counted over the 3,888 homologs with at least 30% sequence identity to CVOMT1 (identical positions divided by aligned query positions) and at least 60% query coverage (aligned query positions divided by the 356 residues of CVOMT1). Of these, 3,597 have a residue at position 264 and 291 have a gap. The counts are unweighted: no clustering or phylogenetic weighting was applied. Substitutions at position 264 were scored with the ESM-1b and ESM-1v zero-shot models[7,8] (wild-type-marginal log-likelihood ratios, averaged over ESM-1b and five ESM-1v models) and with ProteinMPNN-ddG and ThermoMPNN, run on a Boltz-2 model of wild-type CVOMT1 without ligands (complex pLDDT 0.91).[9,10] Per-run values and analyses are listed in Tables S5 to S7.

#### Chromosomal Integrations and De Novo Cultivation

Genomic integrations followed the I-SceI protocol described above. *bsdBCD* was integrated in *P. putida* Δ*pobA* through a Cre/lox landing pad that also carries the T7 RNA polymerase gene and its T7 promoter. *tktA* and *talB* from *E. coli* were integrated under the constitutive tac promoter at the attTn7 site downstream of *glmS*, and FLAG-pomt1 was integrated under the constitutive

J23100 promoter downstream of *PP_5322*. *De novo* cultivations (Figures 4H–K and 5C) were performed in 250 mL Erlenmeyer flasks containing 25 mL M9 minimal medium with 50 mM glucose and a 50% (v/v) dodecane overlay, incubated at 30 °C and 200 rpm. At inoculation (t = 0 h), 2 mM L-arabinose was added to induce expression of the upstream pathway modules: *ubiC* and *bsdBCD* on plasmid pPhenol_4HB (Figure 4H,J) or *menF*, *pchB*, and *sdc* on plasmid pPhenol_SAL (Figure 4I,K), alongside constitutive pomt1 expression from pSEVAb23. Flask cultivations in Figure 5C were sampled periodically across 96 h (4 days).

#### Statistical Analysis

All reported experiments derive from independent biological triplicates unless otherwise noted. Figures show mean values ± standard deviation. Statistical significance was assessed by an unpaired Student's *t*-test. Asterisks in the figures denote *p ≤ 0.05, **p ≤ 0.01, ***p ≤ 0.001 and ****p ≤ 0.0001.

### SUPPLEMENTARY RESULTS

#### Construction of the Selection Strain

To establish a stringent growth-coupled selection, native biosynthetic pathways to cysteine were systematically inactivated. *P. putida* synthesizes cysteine from serine via serine *O*-acetyltransferases, annotated in KEGG as *cysE*, PP_1110, and two putative paralogs, *PP_0228* and *PP_3136*. Simultaneous deletion of annotated *cysE* and PP_1110 failed to disrupt cysteine prototrophy in glucose minimal medium (Figure S2A), indicating functional redundancy from the putative genes. Individual gene deletions revealed their respective roles: deletion of PP_3136 produced no phenotypic defect (Figure S2B), whereas deletion of PP_0228 in the Δ*cysE* Δ*PP_1110* background completely abolished cysteine prototrophy, rendering the strain dependent on supplementation (Figure S2C). These genetic results indicate that *PP_0228* encodes an active serine *O*-acetyltransferase in *P. putida*, whereas *PP_3136* does not sustain detectable flux. To eliminate potential background bypass and fully decouple the synthetic selection module from native metabolism, *PP_3136* was also deleted.

Although transsulfuration of cystathionine can theoretically supply cysteine in *P. putida*, growth assays confirmed that native flux through this pathway is insufficient to rescue auxotrophy (Figure S2D). To eliminate all potential metabolic bypasses and fully uncouple cysteine biosynthesis from central metabolism, we deleted *metX*, which encodes homoserine *O*-acetyltransferase. The resulting double-auxotrophic strain, CM-Aux, exhibited strict dependency on exogenous supplementation of both cysteine and methionine (Figure 2B), establishing a tightly coupled selection chassis.

#### Benchmarking POMT1 against Natural *O*-Methyltransferases

We benchmarked POMT1 against candidate natural *O*-methyltransferases reported to exhibit activity on structurally related aromatics. *O*-methyltransferases frequently display substrate promiscuity toward phenolic compounds.[11] We assembled a panel of candidates using three complementary discovery strategies: literature-curated enzymes with reported phenolic activity (an *O*-methyltransferase (OMT) from *Tetracentron sinense*, a catechol *O*-methyltransferase

(COMT) from *Sesamum indicum*, and a trans-anol *O*-methyltransferase (AIMT) from *Pimpinella anisum*),[12,13] sequence-homology mining with EnzymeMiner for homologs bearing the D264N-equivalent residue, which returned an OMT from *Geodermatophilus obscurus*,[14] and structural similarity search with FoldSeek, which identified an OMT from *Nocardia brasiliensis*.[15] All variants were expressed in the CM-Aux selection strain alongside the C-module and benchmarked against POMT1 and wild-type CVOMT1 at both 1 mM and 0.1 mM phenol (Figure S5).

At 1 mM phenol, AIMT supported cell growth comparable to POMT1 and the *N. brasiliensis* OMT performed similarly to unevolved CVOMT1, while the remaining candidates showed negligible activity (Figure S5D). However, at 0.1 mM phenol, representing the physiologically relevant concentration regime for *de novo* pathway integration, only POMT1 sustained rapid cell growth (Figure S5E). This divergence shows that POMT1 sustains growth at a phenol concentration at which the natural homologs tested do not, which suggests higher activity at low phenol concentration, although kinetic constants were not determined. Notably, the *G. obscurus* enzyme, which was selected for carrying the D264N-equivalent residue, showed negligible activity, so Asn at this position is not sufficient for phenol activity.

**N-Terminal Tags, Expression Context, and Cofactor Supply**

Although POMT1 outperformed CVOMT1, phenol accumulated to 0.98 mM against only 0.16 mM anisole in the glucose-fed 4HB strain (Figure 4J), indicating that its catalytic turnover still constrained pathway throughput. Because CVOMT1 originates from *O. basilicum*, we evaluated N-terminal affinity tags to improve heterologous folding in *P. putida*, drawing on established strategies for heterologous protein solubilization in bacterial hosts.[16] Both a 6×His-tag and a FLAG-tag increased phenol consumption (1.61-fold and 2.93-fold, respectively) and anisole production (2.62-fold and 4.28-fold, respectively) compared with untagged POMT1 (Figure S6A,B), suggesting that folding or solubility limited the functional expression of the untagged enzyme. Because FLAG-POMT1 exhibited the highest conversion efficiency, it was selected for subsequent strain construction. We next examined expression context by evaluating alternative constitutive promoters (*Ptrc* without the *lac* operator, and *PR* without the cI repressor, on pSEVAb23) and the higher-copy pSEVAb24/pRO1600 backbone.[1] Promoter identity made negligible difference on the pSEVAb23 backbone (Figure S6C), but the higher-copy pSEVAb24 backbone reduced both phenol consumption and anisole production (2.05-fold and 1.89-fold, respectively) relative to pSEVAb23, indicating that expression level had passed its optimum and that gene dosage must be balanced rather than maximized.

With expression optimized, we turned to cofactor supply as the next candidate bottleneck. POMT1, like other SAM-dependent methyltransferases, donates a methyl group from SAM and releases SAH, which the native methionine cycle recycles back to SAM via homocysteine and methionine (Figure S6D). Co-expressing the native SAM synthase *metK* with FLAG-POMT1 across varying methionine concentrations (0–2 mM) resulted in no improvement in final anisole titer, in contrast to the benefit of methionine supplementation seen with unevolved CVOMT1 (Figure 1G,H). Conversely, elevated methionine concentrations decreased bioconversion rates in *metK*-overexpressing strains, with phenol consumption falling from 1.82 mM (no methionine) to

1.61, 1.48 and 1.29 mM at 0.5, 1 and 2 mM methionine, respectively, within 24 hours (Figure S7B; corresponding 48 h yields in Figure S6E), consistent with feedback inhibition rather than substrate limitation. All strains nonetheless fully consumed the supplied phenol within 48 hours regardless of methionine or *metK* status (Figure S7). These results indicate that endogenous SAM pools are sufficient to support POMT1 activity, consistent with cofactor availability not being rate-limiting for the terminal methyltransfer reaction. We next investigated whether cofactor *recycling* limits pathway throughput by testing whether accelerated regeneration of SAH could enhance bioconversion. Rather than relying on native *P. putida* genes subject to feedback regulation, we introduced the orthogonal *E. coli mtn-luxS* pathway, which recycles SAH to homocysteine via S-ribosyl-L-homocysteine (Figure S6D), and co-expressed it with FLAG-POMT1 at 2 mM phenol without methionine supplementation. Overexpressing this recycling module produced no improvement in conversion rate or yield (Figure S6F). Together, the *metK* and *mtn-luxS* experiments form a matched pair of negative controls that indicate that neither SAM synthesis nor SAH recycling is rate-limiting under these conditions, narrowing subsequent optimization specifically to the catalytic turnover of POMT1.

#### Stepwise Progression of Titers

Table S8 tracks phenol and anisole titers from the first proof-of-concept conversion through to the complete *de novo* production strain. All values are taken from the corresponding results in the main text and Figures 1, 3, 4 and 5.

**Table S8.** Stepwise improvement in phenol and anisole titers across the engineering campaign, from the first proof-of-concept phenol-to-anisole conversion to the complete *de novo* production strain. n.d., not determined in that experiment.

| Strain / modification | Phenol (mM) | Anisole (mM) | Notes |
|---|---|---|---|
| CVOMT1 + 1 mM phenol (fed, 50% dodecane, 24 h) | 0.049 consumed | 0.038 | Proof of concept; no Met supplement |
| CVOMT1 + 1 mM phenol + 1 mM Met (fed, 50% dodecane, 24 h) | 0.157 consumed | 0.062 | Met supplementation raises conversion |
| CVOMT1 production strain (1 mM phenol, 50% dodecane, 24 h) | 0.30 consumed | 0.06 | Benchmark for POMT1 comparison; separate experiment from rows 1 and 2 |
| POMT1 (D264N) production strain (1 mM phenol, 50% dodecane, 24 h) | 0.39 consumed | 0.14 | 2.3-fold increase over CVOMT1 |
| pAnisole_4HB, 1 mM 4HB (fed, 2 d) | 0.825 | 0.110 | 4HB route, plasmid-based |
| 4HB *de novo* pathway + POMT1, glucose (72 h) | 0.98 | 0.16 | First full *de novo* 4HB-route result |
| PEP-enhanced strain (Δ*pyk* Δ*pykA* Δ*ppc* Δ*pobA* Δ*quiC*), reactor tubes | n.d. | 0.23 (vs. 0.14 for Δ*pobA*) | 64.3% increase from PEP pool expansion |
| + AroF-I P160L (feedback-resistant DAHP synthase) | 1.36 | 0.39 | Largest single-step gain |
| + TrpE P290S (reduced tryptophan flux) | 1.70 | 0.56 | Plasmid-based FLAG-POMT1 |
| Chromosomal FLAG-POMT1 (same background) | 1.87 | 0.20 | Quantifies gene-dosage dependence |
| Final *de novo* strain, shake flask, glucose (48 h) | 1.66 | 0.57 | Complete optimized strain |

**SUPPLEMENTARY FIGURES**

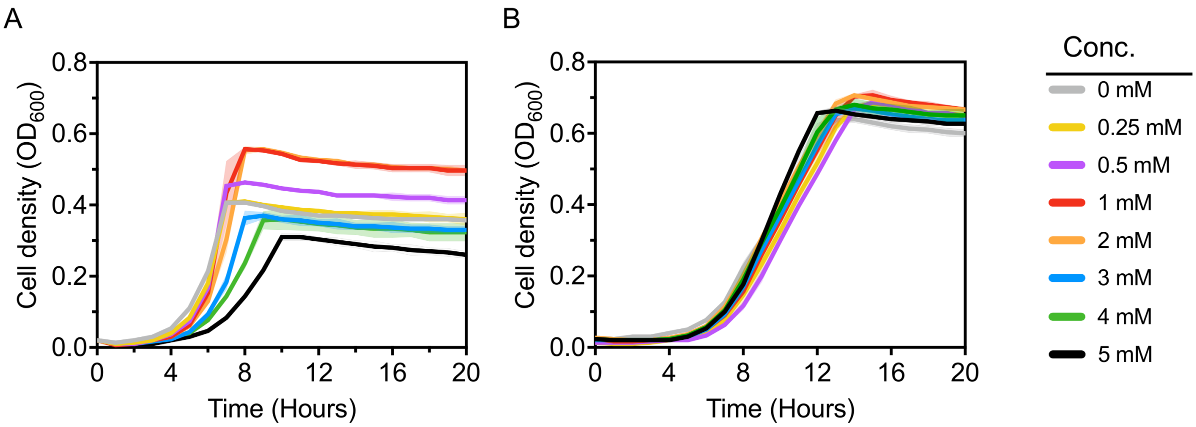


**Figure S1.** Toxicity analyses. (A) Growth curves of *P. putida* KT2440 subjected to different phenol concentrations. (B) Growth curves of *P. putida* KT2440 subjected to different anisole concentrations. Growth curves represent the mean value ± SD from three independent experiments.

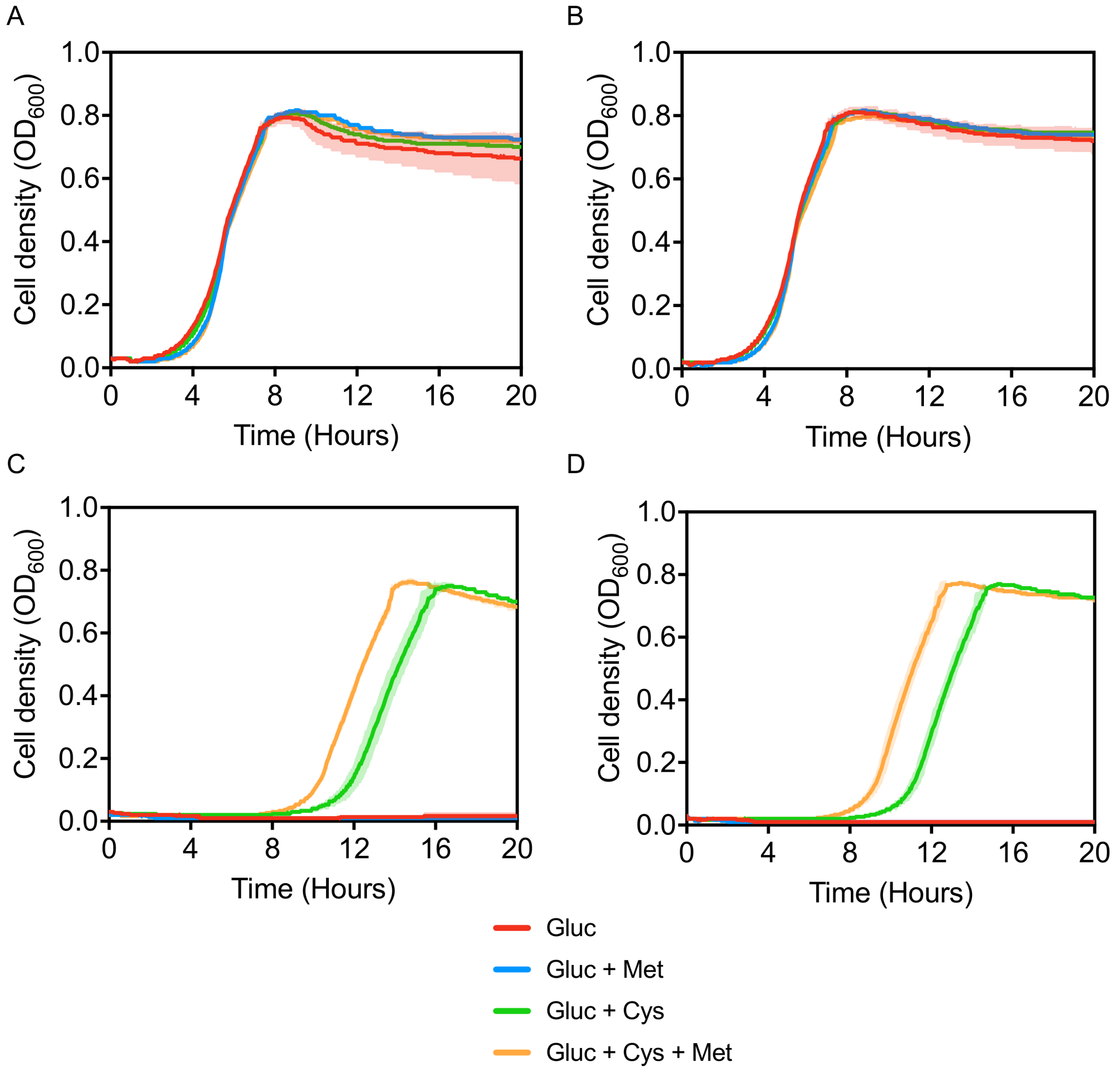


**Figure S2.** Construction and characterization of a cysteine auxotroph. (A) *P. putida* Δ*cysE* Δ*PP_1110*. (B) *P. putida* Δ*cysE* Δ*PP_1110* Δ*PP_3136*. (C) *P. putida* Δ*cysE* Δ*PP_1110* Δ*PP_0228*. (D) *P. putida* Δ*cysE* Δ*PP_1110* Δ*PP_0228* Δ*PP_3136*. Abbreviations: (Gluc) glucose, (Cys) cysteine, (Met) methionine. Growth curves represent the mean value ± SD from three independent experiments.

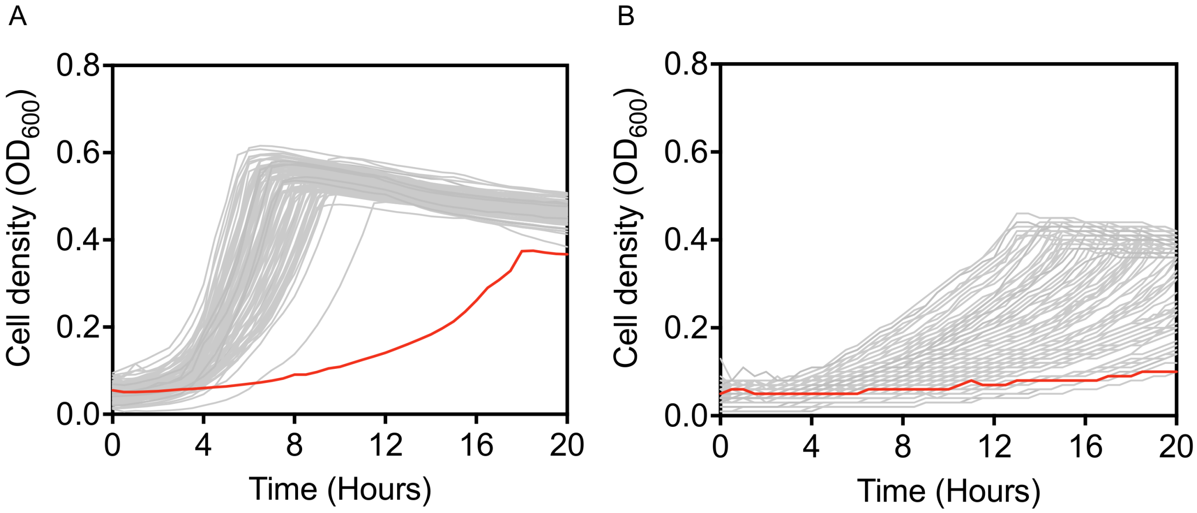


**Figure S3.** Growth phenotyping of CVOMT1 library isolates at 0.1 and 0.01 mM phenol. (A) Growth curves of CM-Aux with the implemented C- and the mutated A-module at a phenol concentration of 0.1 mM. (B) Growth curves of CM-Aux with the implemented C- and the mutated A-module at a phenol concentration of 0.01 mM. Every line represents a single isolate. Growth-coupled selection with the unevolved CVOMT enzyme is indicated in red.

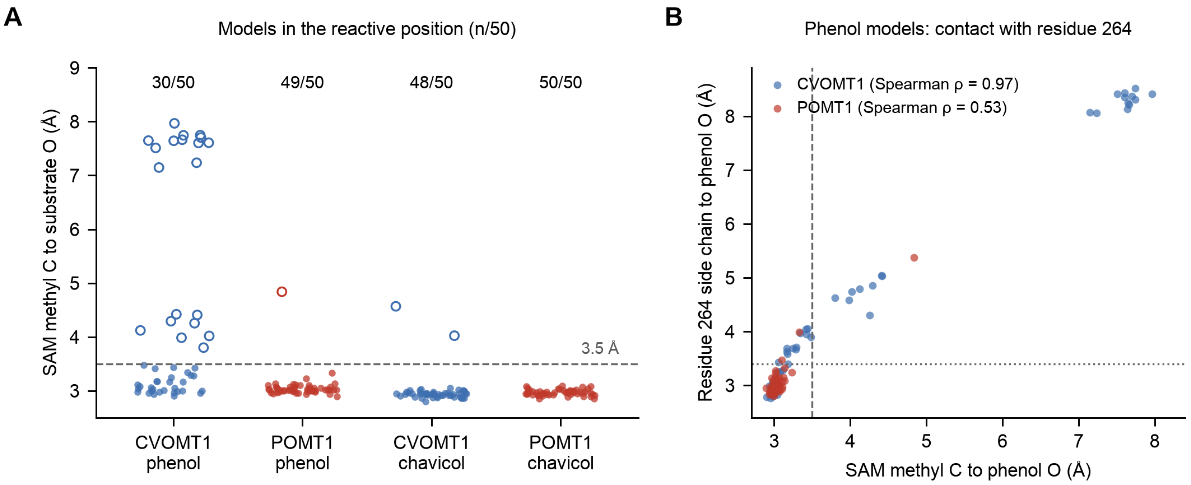


**Figure S4.** Predicted placement of phenol and chavicol in CVOMT1 and POMT1. (A) Distance between the SAM methyl carbon and the substrate oxygen in each of 50 Boltz-2 models per system (two independent runs of 25 samples, one sequence alignment shared by both enzymes). Filled symbols are models within 3.5 Å (dashed line), the cut-off used to call a model reactive, and open symbols are models outside it. The numbers above the panel give reactive models per 50. (B) Distance from the phenol oxygen to the nearest side-chain oxygen or nitrogen of residue 264, plotted against the SAM methyl distance for the phenol models (dotted line, 3.4 Å contact cut-off), with the Spearman correlation for each enzyme. Counts are given in Table S6. All models are predictions.

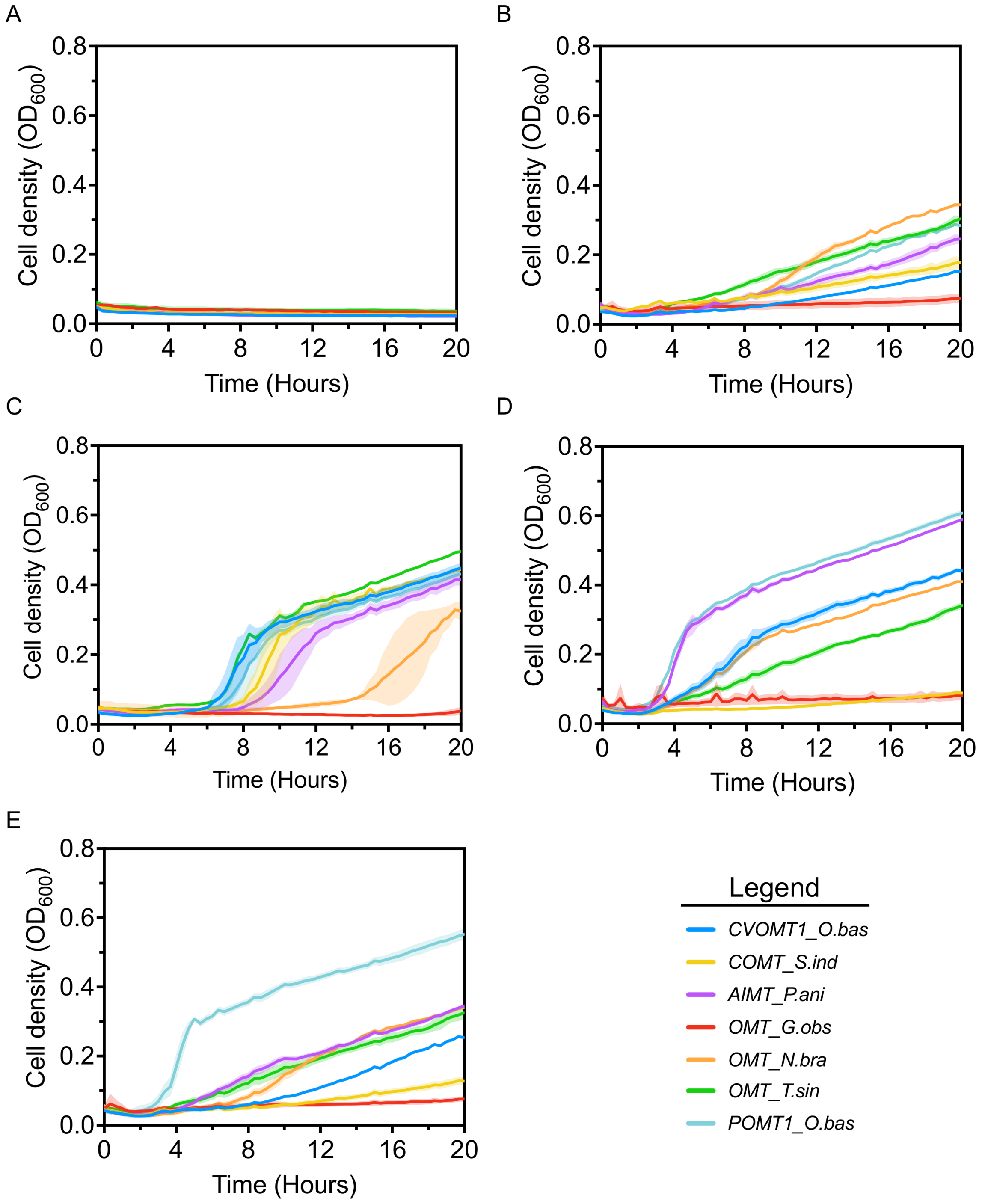


**Figure S5.** Screening of alternative phenol *O*-methyltransferases for anisole production. (A) 10 mM glucose. (B) 10 mM glucose + 1 mM methionine. (C) 10 mM glucose + 1 mM methionine + 1 mM cysteine. (D) 10 mM glucose + 1 mM methionine + 1 mM phenol. (E) 10 mM glucose + 1 mM methionine + 0.1 mM phenol. Abbreviations: (CVOMT1) chavicol *O*-methyltransferase 1, (COMT) catechol *O*-methyltransferase, (AIMT) trans-anol *O*-methyltransferase, (OMT) *O*-methyltransferase, (POMT1) phenol *O*-methyltransferase 1, (O.bas) *Ocimum basilicum*, (S.ind)

*Sesamum indicum*, (P.ani) *Pimpinella anisum*, (G.obs) *Geodermatophilus obscurus*, (N.bra) *Nocardia brasiliensis*, (T.sin) *Tetracentron sinense*.

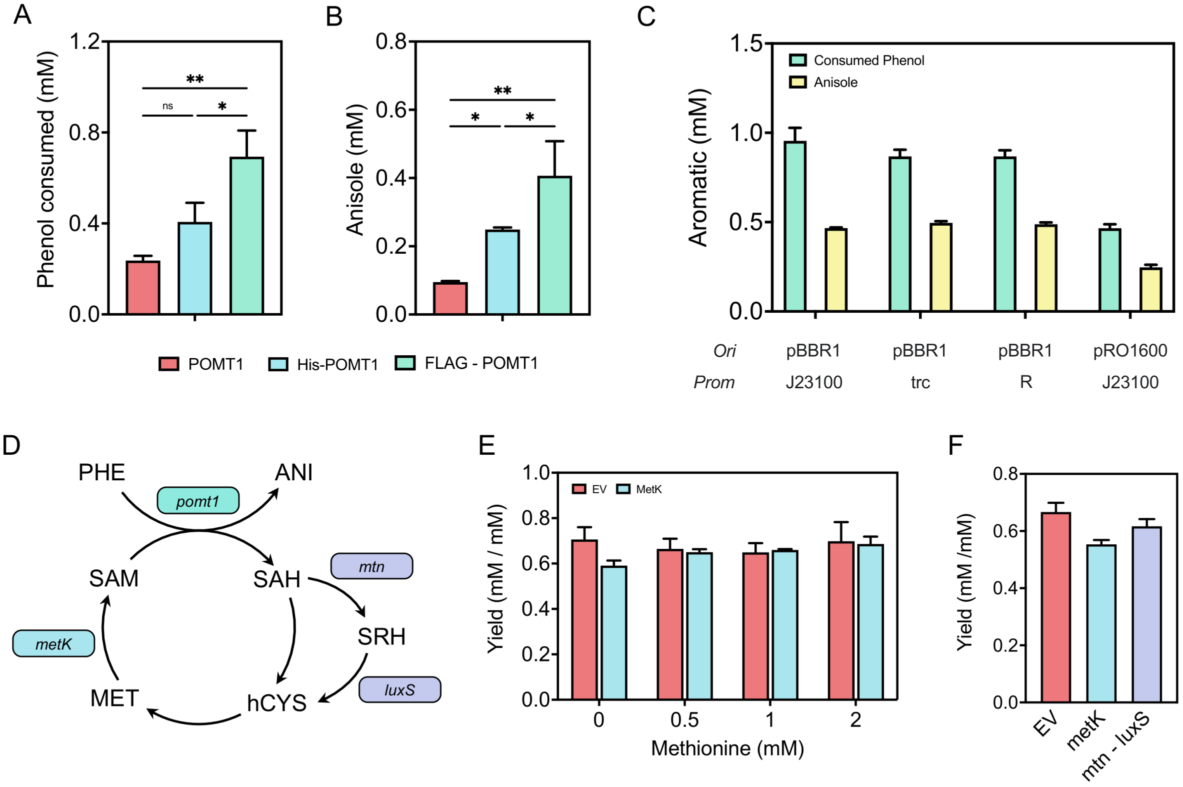


**Figure S6.** Optimization of anisole biosynthesis. (A) Phenol consumption by POMT1 with and without solubility tags. (B) Anisole production by POMT1 with and without solubility tags. (C) Phenol consumption and anisole production by FLAG-POMT1 under different promoters and/or origins of replication. (D) Schematic of the methionine cycle in *P. putida*. (E) Anisole yield (mol anisole produced per mol initial phenol supplied, 2 mM) at 48 h across different methionine concentrations with and without *metK* overexpression. (F) Anisole yield at 48 h upon overexpression of key methionine-cycle enzymes (FLAG-POMT1 alone or co-expressed with the E. coli mtn-luxS recycling cassette). Abbreviations: (MET) methionine, (SAM) S-adenosylmethionine, (SAH) S-adenosyl-L-homocysteine, (hCYS) homocysteine, (SRH) S-ribosyl-L-homocysteine, (PHE) phenol, (ANI) anisole, (EV) empty vector. Genes: *metK* (methionine adenosyltransferase), *pomt1* (phenol *O*-methyltransferase), *mtn* (S-adenosylhomocysteine nucleosidase), *luxS* (S-ribosylhomocysteine lyase). Data points represent the mean value ± SD from three independent experiments.

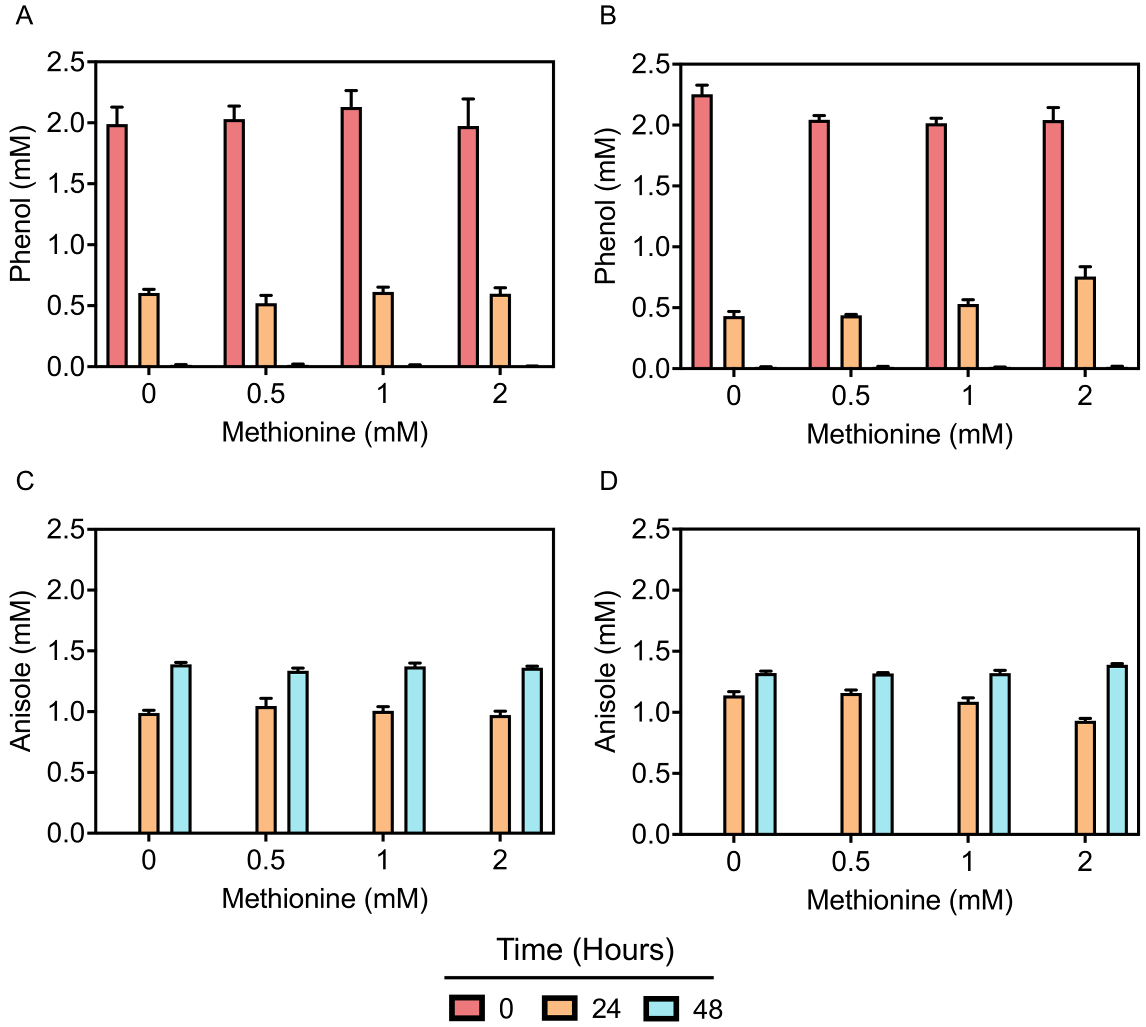


**Figure S7.** Effect of methionine supplementation on whole-cell bioconversion. All strains express FLAG-POMT1 (pSEVAb23) and carry either the empty vector control (pSEVAb62; A, C) or the SAM synthase overexpression plasmid pMetK (pSEVAb62-metK; B, D). Bioconversions were supplied with 2 mM phenol and supplemented with 0, 0.5, 1.0, or 2.0 mM L-methionine in M9 minimal medium with a 50% (v/v) dodecane overlay. (A) Phenol consumption in the control strain. (B) Phenol consumption in the metK-overexpressing strain. (C) Anisole production in the control strain. (D) Anisole production in the metK-overexpressing strain. Data points represent the mean ± SD from three independent biological replicates.
